## Supplementary Material for "Assessing the performance of protein regression models"

### SUPPLEMENTARY MATERIAL
### Assessing the performance of protein regression models

Richard Michael*[1], Jacob Kæstel-Hansen*[2], Peter Mørch Groth[1,3], Simon Bartels[1], Jesper Salomon[3], Pengfei Tian[3], Nikos S. Hatzakis[2], Wouter Boomsma[1]

[1]Department of Computer Science, University of Copenhagen

[2]Department of Chemistry, University of Copenhagen

[3]Bioinformatics & Design, Enzyme Research, Novozymes A/S

*These authors contributed equally to this work.

#### 1 DATA

Supplementary Table 1: Overview of the Data.

| DATASET | REF. | EFFECT | STRESS TYPE |
|---|---|---|---|
| β-Lactamase | (43) | growth | CHEMICAL |
| Ubiquitin | (44) | growth | CHEMICAL |
| Calmodulin | (45) | growth | CHEMICAL |
| TIM-Barrel | (46) | growth | THERMAL |
| BRCA1 | (47) | growth | CHEMICAL |
| T2-MTH | (48) | growth | CHEMICAL/GENETIC |
| ParD-antitoxin | (36) | growth | CHEMICAL |

For each protein we have experimental observations from deep mutational scans (DMS), which record growth under chemical stress - β-Lactamase, Ubiquitin, BRCA1, ParD-antitoxin, and thermal TIM-Barrel (see table 1 cf. supplementary material in[(8)]).

#### 2 COMPUTING INFRASTRUCTURE

All experiments were initially run on an Apple M1 Pro (using the TensorFlow Metal support), and subsequently a compute cluster with Intel Xeon 6248 CPUs, running a Linux kernel 4.18.0-425.3.1.e18.x86_64 and NVIDIA Quadro RTX 6000 (Driver Vers. 525.60.13), NVIDIA TITAN X, using CUDA 11.6 , 11.8, and 12.0 . Representations were obtained using the aforementioned compute cluster with respective specifications.

### 3 ADDITIONAL RESULTS

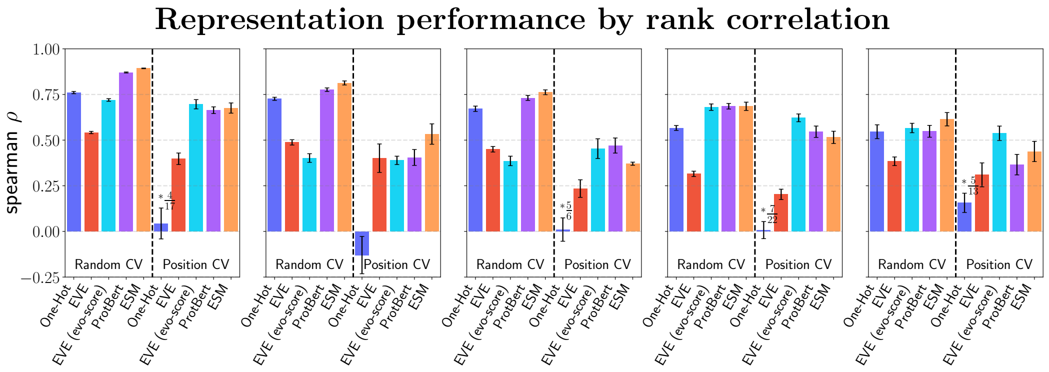


Supplementary Figure 1: Representation comparison of GP (Matérn$\frac{5}{2}$) by spearman rank-correlation. The annotations indicate the fraction of computable results: i.e. if the predictions are constant as is the case for some data-splits with ONE-HOT input the *spearman* $\rho$ cannot be computed and only the results of splits are presented, which are not constant.

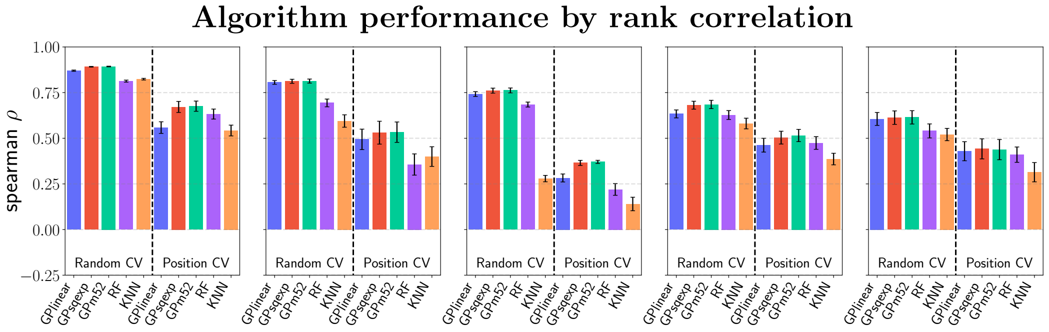


Supplementary Figure 2: Algorithm comparison on ESM-1B representation by rank-correlation.

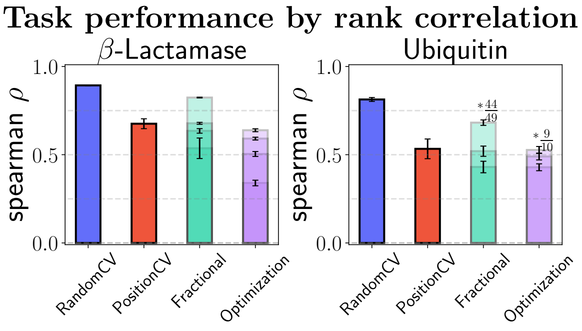


Supplementary Figure 3: Task comparison of GP (Matérn$\frac{5}{2}$ on ESM-1B) by rank-correlation. The annotations indicate the fraction of computable results.

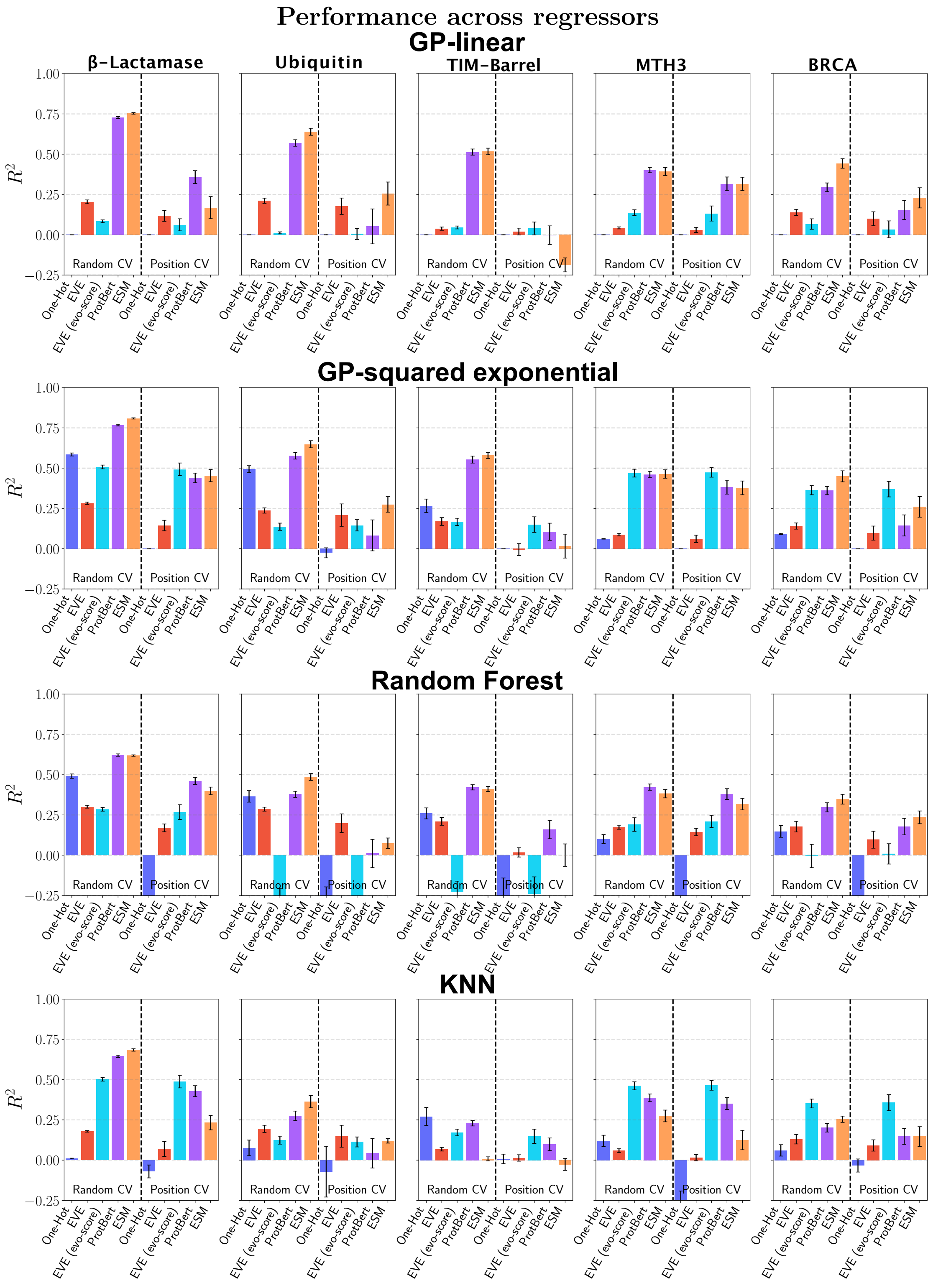


Supplementary Figure 4: Comparison of the performance ($R^2$) of individual regressors (rows) by embeddings for splitting at random and by position.

### 4 CONSIDERATIONS ON THE INDEPENDENCE OF PROTEIN VARIANTS

In addition to the described domains for protein variants, their distributions and properties the following situation arises.: Depending on the correlation of the positions and type of mutation multiple variants

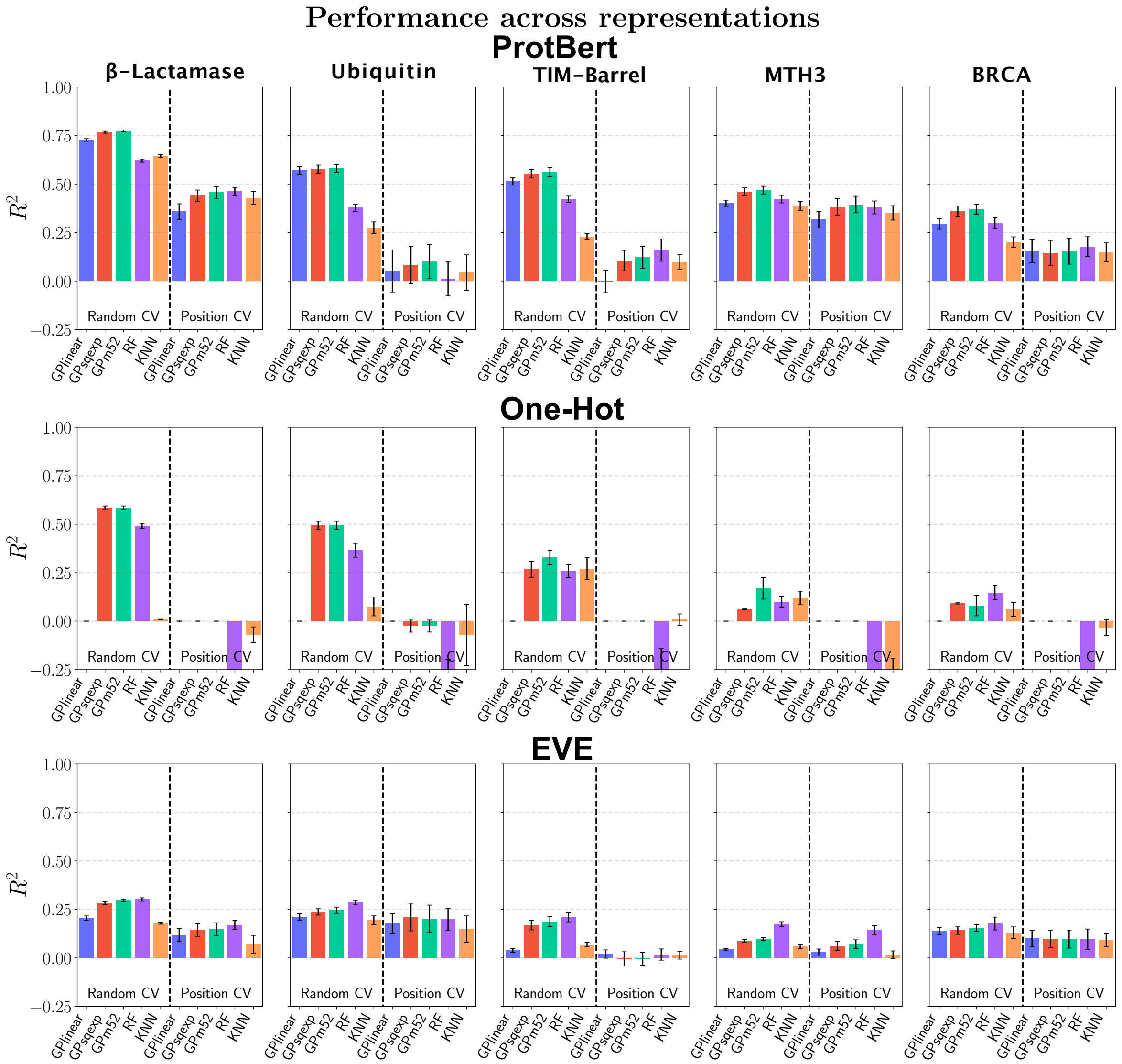


Supplementary Figure 5: Individual representation (rows) performance ($R^2$) across the available regressors (x-axis).

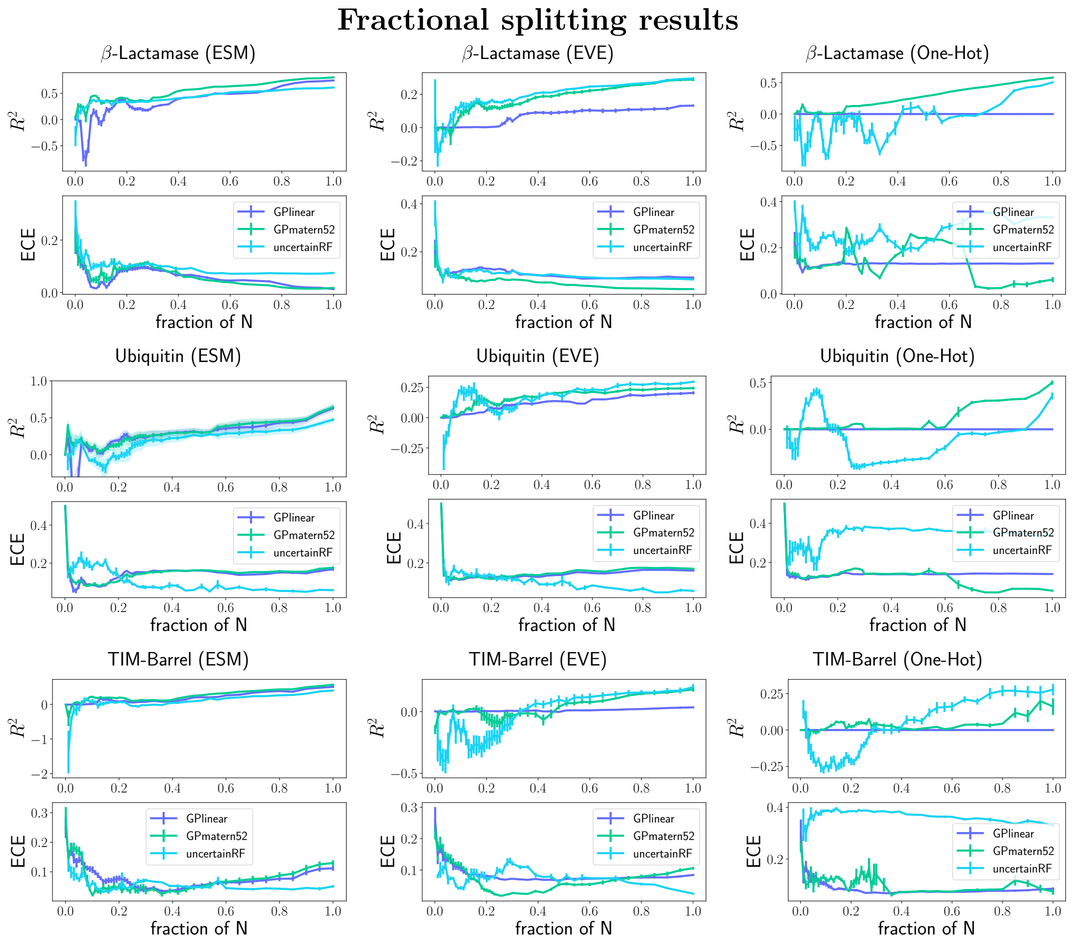


Supplementary Figure 6: Individual performance ($R^2$) and deviation from calibration (ECE) across fractions of available training data (splitting protocol, 5-fold CV per fraction), for $\beta$-Lactamase, Ubiquitin, TIM-Barrel, three different regressors (GP linear, GP squared-exponential, and Random Forest with uncertainty estimate). The bars indicate std.err. across CV splits.
Note that for the $\beta$-Lactamase *uncertain RF* regressor experiments the optimal parameters from the random CV protocols were used for the second half of the data-fraction experiments across all representation, specifically splits {0.36, 0.39, ... 0.95, 1.0} for computational feasibility. This only applies for the RF experiments.

may or may *not* be considered *independent*. Let $m_{\text{A}}$ and $m_{\text{B}}$ be two mutations on a protein variant sequence $S$. Then $m_{\text{A}} \perp\!\!\!\perp m_{\text{B}}$ on $S$, iff

1. $m_{\text{A}}$ is sufficiently distant to $m_{\text{B}}$, such that the residue properties do not impose on one another given the secondary and tertiary structure of the sequence, e.g. a distance around each residue of r$>$ 5Å,

2. the evolutionary co-occurrence of the residues is not significant.

The latter point can be computed by the sequence's evolutionary statistical energy $E \in [0, 1]$, and a dependency threshold $t \in [0, 1]$ then

$$\Delta E(S \setminus m_{\text{A}}, S \setminus m_{\text{B}}) < t \tag{10}$$

(see (3) in [6]).
In contrast, two mutations on one sequence which are in close proximity and or have a high covariate dependency may be treated as one unit for downstream evaluation purposes.

#### 5 MULTIMODALITY OF OBSERVATIONS

Sequences can be categorized into and *high* fitness ($Y^{(kM)^+} := \{\forall y \in Y^{(kM)} : y > t\}$) and *low* fitness ($Y^{(kM)^-} := \{Y^{(kM)} \backslash Y^{(kM)^+}\}$) relative to the initial wild-type observation or a threshold ($t \in \mathbb{R}$) of interest. Generally, for such data, we observe significantly more *low* variates: $|Y^{(kM)^+}| \ll |Y^{(kM)^-}|$ when higher order variants are considered One can look at the modality in the observation distribution by a WT threshold (this applies particularly to Fig. 6). To investigate this effect two scenarios are possible.: Either, we filter the available data to adapt the domain and task respective a functional threshold - selecting $\{(X_i, y_i)|y_i \geq \mathrm{t}\}$, where $t$ is an expected threshold, e.g. the WT performance. Alternatively, we use all available data during training, but only measure performance on a subset; thus filter given the threshold mentioned above.

#### 6 $\chi^2$ CALIBRATION

When considering the (reduced) $\chi^2$ statistic for the goodness of fit, the normalization constant can additionally be extended to account for the degrees of freedom of the regressor. This is well defined for the Random Forest [57], however less so for non-parametric models such as the Gaussian Process regressors. Therefore the metric in its presented equation encapsulates also the inherent model complexity.
To compute the metric in practice we make the assumption of independence between predictions under the model, specifically $\{\hat{\sigma}^2\}_{1..N}$ such that we can build a covariance matrix from the diagonal of the predictive variances and solve by cholesky factorization - as done in [55]. This is equivalent to the previously stated sum of residuals (normalized by the predictive variances) under the previously stated assumptions.

#### 7 FRACTIONAL SPLITTING

The performance of the Random Forest and GP (Matérn$\frac{5}{2}$) shows a monotonic increase over the fractions (for the majority of data-sets and representations); while all regressors show performance fluctuations with low sequence numbers. All methods steadily improve after approx. 30% available training data. All models appear sufficiently well calibrated. Only on the ONE-HOT representation the ECE values of the GPs are significantly lower compared to the RF. The performance suggests GP-Matérn and RF to perform well, both on individual fractions. We note that the performance of all methods on the ESM-1B representation is comparable to that of the shown PROTBERT model. In contrast to the language models, the regressors'performance deteriorate on the ONE-HOT embedding of $\beta$-LACTAMASE.

### 8 OPTIMIZATION RESULTS

After an initial random selection, we add the next best candidate sequence by its highest expected reward from the predictions of the regressor, and then retrain the model, with the new observation in the data-set. We have two reference baselines: 1) selecting sequences at random, and 2) a static protocol, which iterates over a fixed score-ranking proposal of an unsupervised model.[1] At each step we report the function value of the currently selected best sequence candidate, the average across all observations, and the cumulative regret (SI Fig.15)). The results on the ProtBert representation show that the GP based models find the optimal candidate after the fewest observations (see Fig.15). The static, unsupervised baseline demonstrates strong performance in turn of the obtained mean and regret scores, but ultimately uses more iterations to find the optimal solution. The reason for this behavior is that the EVE model orders by evo-score, such that low value observations are placed within the first 500 iterations, while the regressors do not perform such an explicit ordering. Instead, the optimization method iteratively scores the variants by its *acquisition function* and updates the scores at each iteration. Though the Random Forest method has previously shown comparatively high accuracy, the absolute best value is not found within the allotted budget. The predictive uncertainties for the random forest stay constant during extrapolation respective the expected ensemble variance of the closest known value. This stands in contrast to the GP uncertainties, which increase for extrapolation predictions. This leads to less exploration of sequences by the RF, which prohibits us to find the best candidate. On the other hand, the properties of the GPs allow for exploration in regions of higher uncertainties. We note that the performance is similar between ProtBert and the ESM-1b representation, with the best observations being found after the evo-scores (over 300 steps) instead (see SI Fig.15). These additional results show that we cannot quantify the exact expected optimization performance, given a cumulative metric and calibration, but can only obtain an approximate order for the assessed methods and are dependent on the representation.

[1] In our case this is the evolutionary scoring of the EVEmodel, which is the relative loss-bound across 2000 samples.

Supplementary Table 2: **Train- Test-Set sizes Random CV and Positional CV.**
Overview of the sizes of the data sets for [*training* ; *test*] for the initial benchmarking protocol splits (rows) - standard 10-fold Cross Validation and splitting by positions. Note that due to the difference in length between the protein sequences there is a different amount of splits available for each. The representation column accounts for the ONE-HOT, PROTBERT, ESM-1B, EVE embeddings, while eve-density is a subset of EVE and for some proteins there are less sequences available, compared to the total dataset.

| | | $\beta$-LACTAMASE | | UBIQUITIN | | TIM-BARREL | | T2-MTH | | BRCA1 | |
|---|---|---|---|---|---|---|---|---|---|---|---|
| | | representations | eve_density | representations | eve_density | representations | eve_density | representations | eve_density | representations | eve_density |
| **Random CV** | **0** | [4309;479] | [4309;479] | [1074;120] | [1043;116] | [1367;152] | [1367;152] | [1547;172] | [1547;172] | [934;104] | [934;104] |
| | **1** | [4309;479] | [4309;479] | [1074;120] | [1043;116] | [1367;152] | [1367;152] | [1547;172] | [1547;172] | [934;104] | [934;104] |
| | **2** | [4309;479] | [4309;479] | [1074;120] | [1043;116] | [1367;152] | [1367;152] | [1547;172] | [1547;172] | [934;104] | [934;104] |
| | **3** | [4309;479] | [4309;479] | [1074;120] | [1043;116] | [1367;152] | [1367;152] | [1547;172] | [1547;172] | [934;104] | [934;104] |
| | **4** | [4309;479] | [4309;479] | [1075;119] | [1043;116] | [1367;152] | [1367;152] | [1547;172] | [1547;172] | [934;104] | [934;104] |
| | **5** | [4309;479] | [4309;479] | [1075;119] | [1043;116] | [1367;152] | [1367;152] | [1547;172] | [1547;172] | [934;104] | [934;104] |
| | **6** | [4309;479] | [4309;479] | [1075;119] | [1043;116] | [1367;152] | [1367;152] | [1547;172] | [1547;172] | [934;104] | [934;104] |
| | **7** | [4309;479] | [4309;479] | [1075;119] | [1043;116] | [1367;152] | [1367;152] | [1547;172] | [1547;172] | [934;104] | [934;104] |
| | **8** | [4310;478] | [4310;478] | [1075;119] | [1043;116] | [1367;152] | [1367;152] | [1547;172] | [1547;172] | [935;103] | [935;103] |
| | **9** | [4310;478] | [4310;478] | [1075;119] | [1044;115] | [1368;151] | [1368;151] | [1548;171] | [1548;171] | [935;103] | [935;103] |
| **Position CV p=15** | **0** | [4427;285] | [4427;285] | [898;220] | [863;220] | [1159;284] | [1159;284] | [1620;79] | [1620;79] | [933;82] | [933;82] |
| | **1** | [4351;285] | [4351;285] | [797;258] | [762;258] | [1082;285] | [1082;285] | [1597;79] | [1597;79] | [912;82] | [912;82] |
| | **2** | [4351;285] | [4351;285] | [829;260] | [794;260] | [1082;285] | [1082;285] | [1596;81] | [1596;81] | [904;87] | [904;87] |
| | **3** | [4351;285] | [4351;285] | [824;230] | [789;230] | [1082;285] | [1082;285] | [1591;83] | [1591;83] | [914;76] | [914;76] |
| | **4** | [4351;285] | [4351;285] | [900;226] | [900;191] | [1082;285] | [1082;285] | [1588;85] | [1588;85] | [982;36] | [982;36] |
| | **5** | [4351;285] | [4351;285] | | | [1348;95] | [1348;95] | [1596;82] | [1596;82] | [911;90] | [911;90] |
| | **6** | [4351;285] | [4351;285] | | | | | [1595;81] | [1595;81] | [912;78] | [912;78] |
| | **7** | [4351;285] | [4351;285] | | | | | [1588;84] | [1588;84] | [912;83] | [912;83] |
| | **8** | [4484;228] | [4484;228] | | | | | [1595;82] | [1595;82] | [905;86] | [905;86] |
| | **9** | [4351;285] | [4351;285] | | | | | [1599;77] | [1599;77] | [905;87] | [905;87] |
| | **10** | [4351;285] | [4351;285] | | | | | [1599;77] | [1599;77] | [905;84] | [905;84] |
| | **11** | [4351;285] | [4351;285] | | | | | [1589;83] | [1589;83] | [910;84] | [910;84] |
| | **12** | [4351;285] | [4351;285] | | | | | [1597;79] | [1597;79] | [912;83] | [912;83] |
| | **13** | [4351;285] | [4351;285] | | | | | [1600;80] | [1600;80] | | |
| | **14** | [4351;285] | [4351;285] | | | | | [1679;18] | [1679;18] | | |
| | **15** | [4351;285] | [4351;285] | | | | | [1592;83] | [1592;83] | | |
| | **16** | [4351;285] | [4351;285] | | | | | [1602;78] | [1602;78] | | |
| | **17** | | | | | | | [1599;77] | [1599;77] | | |
| | **18** | | | | | | | [1593;82] | [1593;82] | | |
| | **19** | | | | | | | [1592;82] | [1592;82] | | |
| | **20** | | | | | | | [1591;82] | [1591;82] | | |
| | **21** | | | | | | | [1592;85] | [1592;85] | | |

Supplementary Table 3: **Train- Test-Set sizes mutational CV.**
Overview of the sizes of the data sets for [*training* ; *test*] for each mutational split (rows). Intra-domain assessment ($1 \to 1$, $2 \to 2$, $3 \to 3$) is standard 5-fold cross-validation. Across domains (extrapolation) of mutations compared to the *WT* are $1 \to 2$, $\leq 2 \to 3$, $\leq 3 \to 4$.

| | | PARD-ANTITOXIN |
|---|---|---|
| **BioSplitter1_1** | **0** | [31;7] |
| | **1** | [31;7] |
| | **2** | [31;7] |
| | **3** | [31;7] |
| | **4** | [31;7] |
| **BioSplitter1_2** | **0** | [38;499] |
| **BioSplitter2_2** | **0** | [438;99] |
| | **1** | [438;99] |
| | **2** | [438;99] |
| | **3** | [438;99] |
| | **4** | [438;99] |
| **BioSplitter2_3** | **0** | [537;2798] |
| **BioSplitter3_3** | **0** | [2776;559] |
| | **1** | [2776;559] |
| | **2** | [2776;559] |
| | **3** | [2776;559] |
| | **4** | [2776;559] |
| **BioSplitter3_4** | **0** | [3335;5858] |

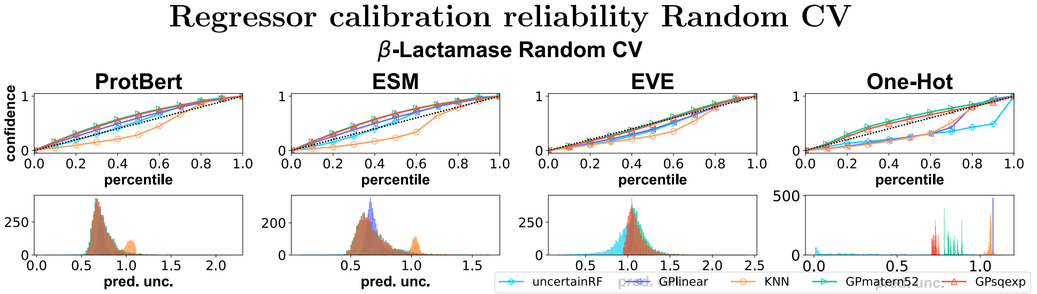


(a) Calibration by reliability curves (top) of $\beta$-Lactamase split at random and histogram of predictive uncertainties (bottom).

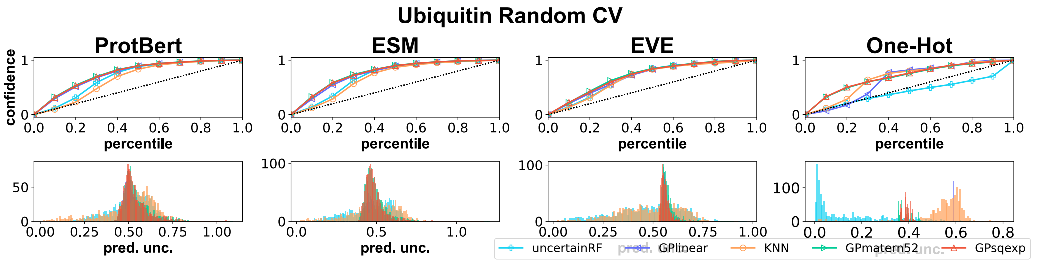


(b) Calibration by reliability curves (top) of Ubiquitin for splitting at random and histogram of predictive uncertainties (bottom).

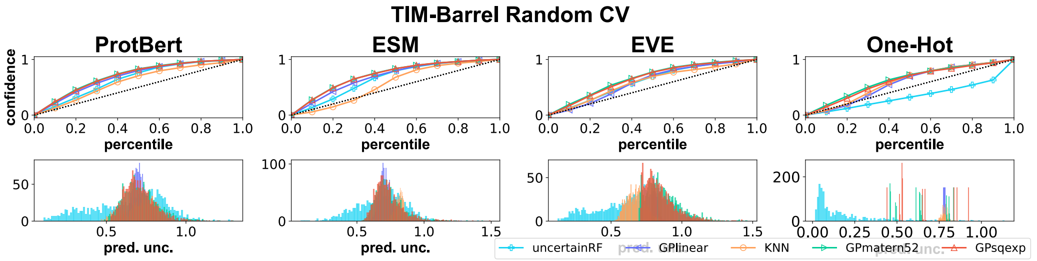


(c) Calibration by reliability curves (top) of TIM-Barrel for splitting at random and histogram of predictive uncertainties (bottom).

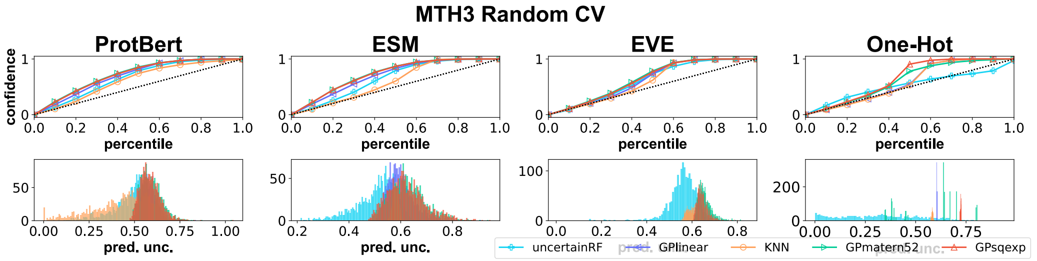


(d) Calibration by reliability curves (top) of T2-MTH for splitting at random and histogram of predictive uncertainties (bottom).

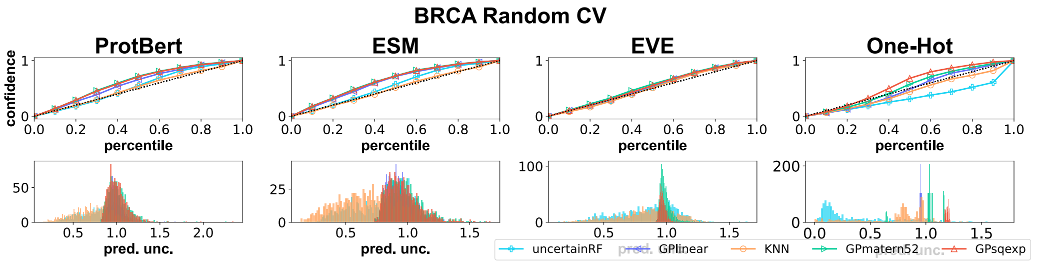


(e) Calibration by reliability curves (top) of BRCA1 for splitting at random and histogram of predictive uncertainties (bottom).

Supplementary Figure 7: Calibration of regressors by reliability curves for splitting by Random CV.

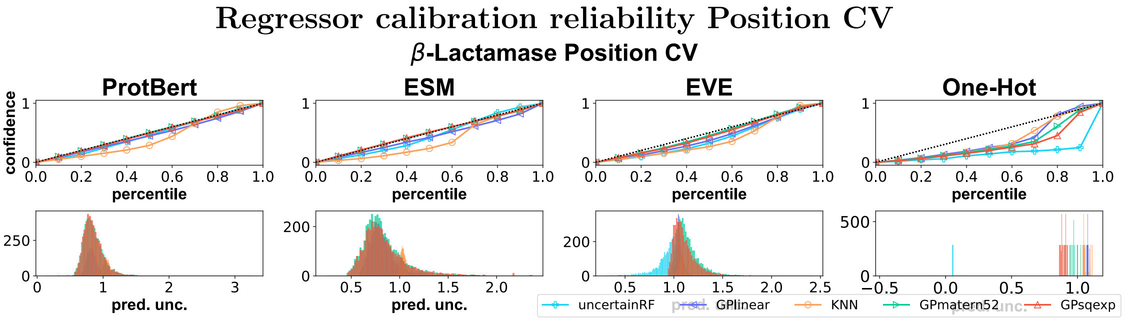


(a) Calibration by reliability curves (top) of $\beta$-LACTAMASE for splitting by position and histogram of predictive uncertainties (bottom).

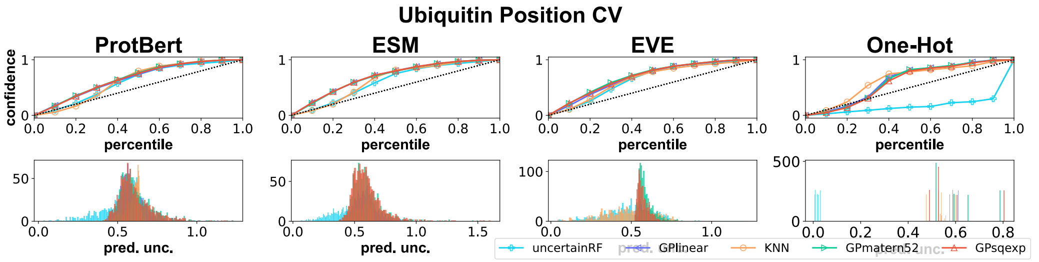


(b) Calibration by reliability curves (top) of UBIQUITIN for splitting by position and histogram of predictive uncertainties (bottom).

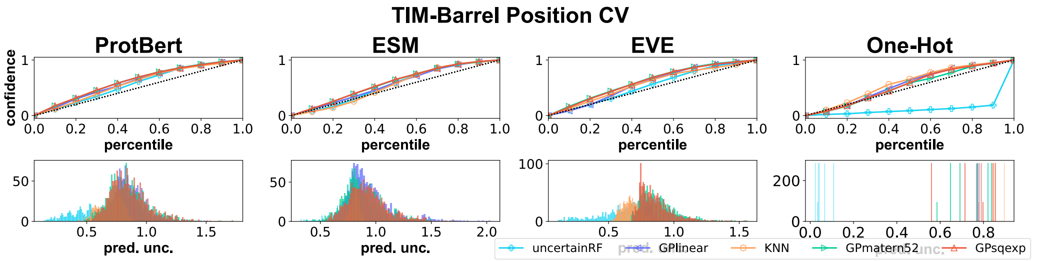


(c) Calibration by reliability curves (top) of TIM-BARREL for splitting by position and histogram of predictive uncertainties (bottom).

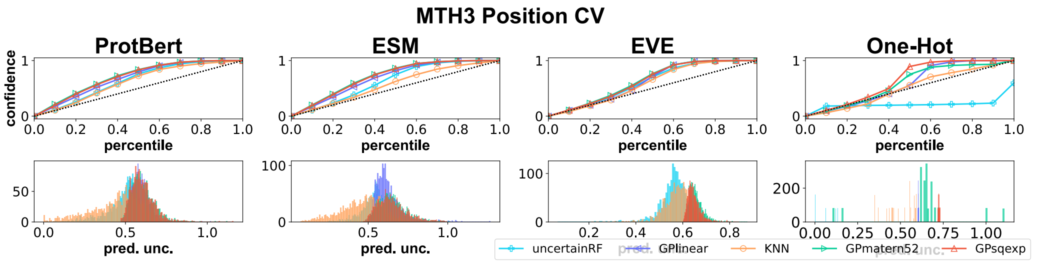


(d) Calibration by reliability curves (top) of T2-MTH for splitting by position and histogram of predictive uncertainties (bottom).

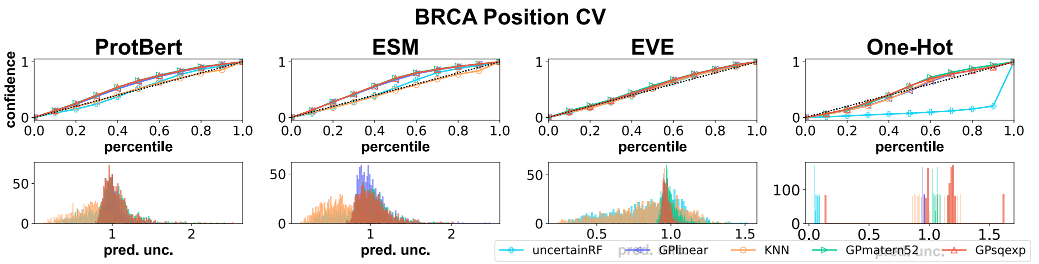


(e) Calibration by reliability curves (top) of BRCA1 for splitting by position and histogram of predictive uncertainties (bottom).

Supplementary Figure 8: Calibration of regressors by reliability curves for splitting by position.

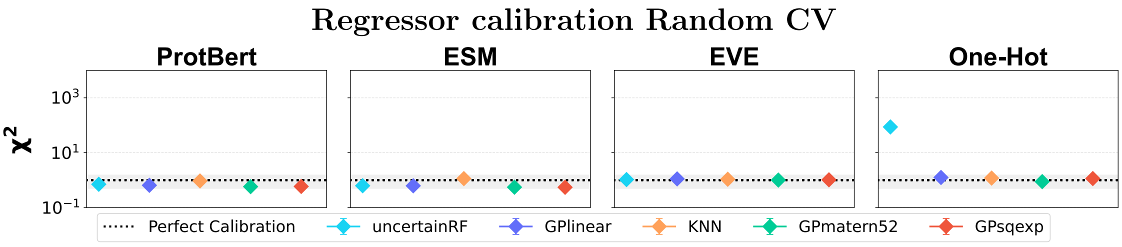


(a) $\chi^2$ calibration of regressors on $\beta$-Lactamase when splitting at random.

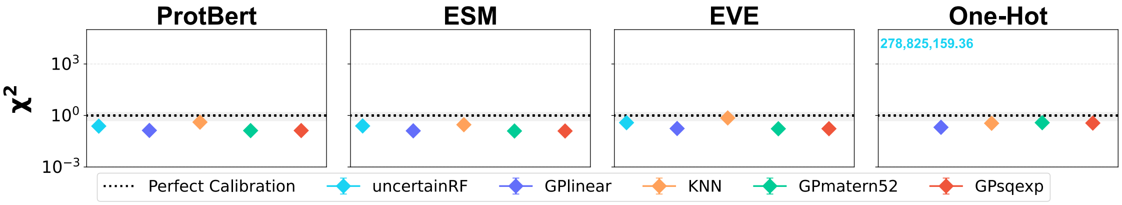


(b) $\chi^2$ calibration of regressors on Ubiquitin when splitting at random.

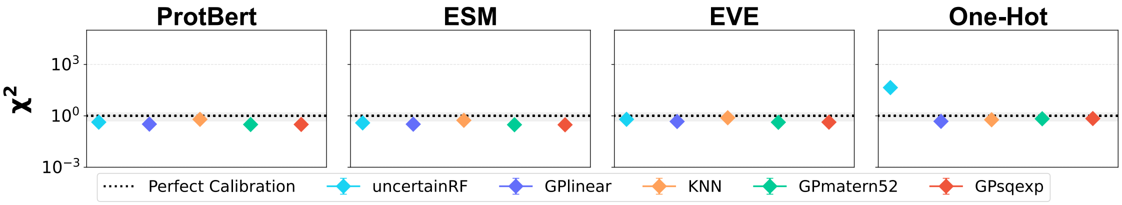


(c) $\chi^2$ calibration of regressors on TIM-Barrel when splitting at random.

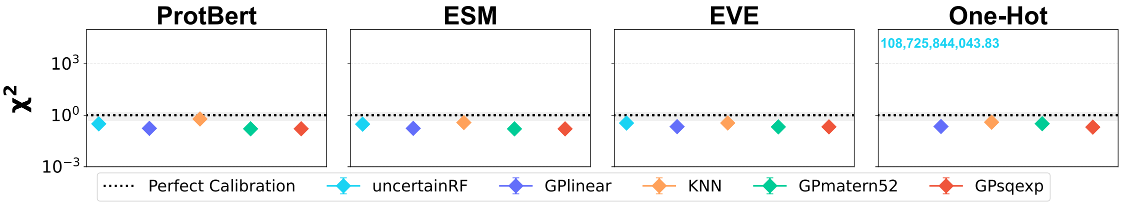


(d) $\chi^2$ calibration of regressors on T2-MTH when splitting at random.

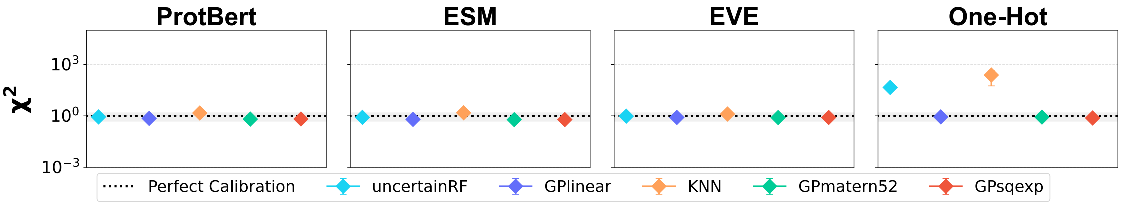


(e) $\chi^2$ calibration of regressors on BRCA1 when splitting at random.

Supplementary Figure 9: Calibration of regressors by (reduced) $\chi^2$ for splitting at random.

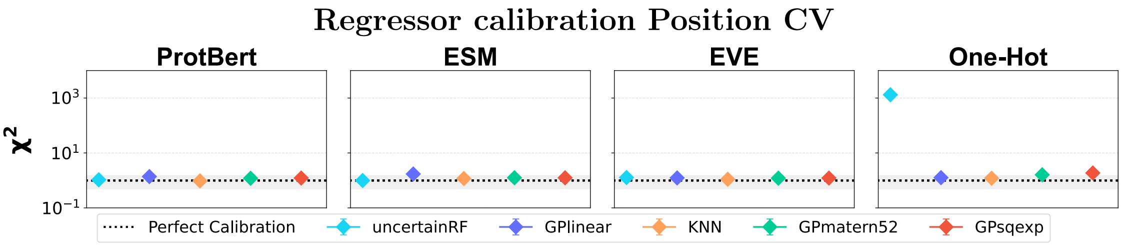


(a) $\chi^2$ calibration of regressors on $\beta$-Lactamase when splitting by position.

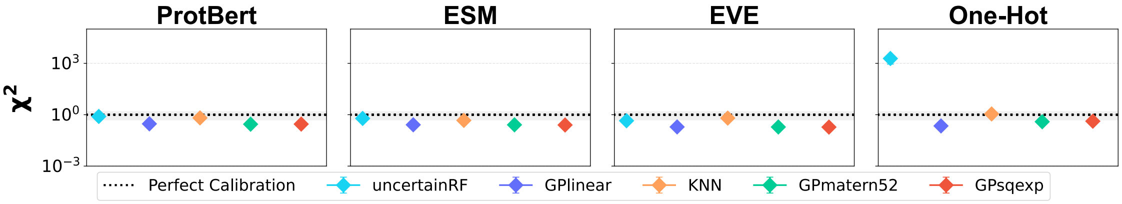


(b) $\chi^2$ calibration of regressors on Ubiquitin when splitting by position.

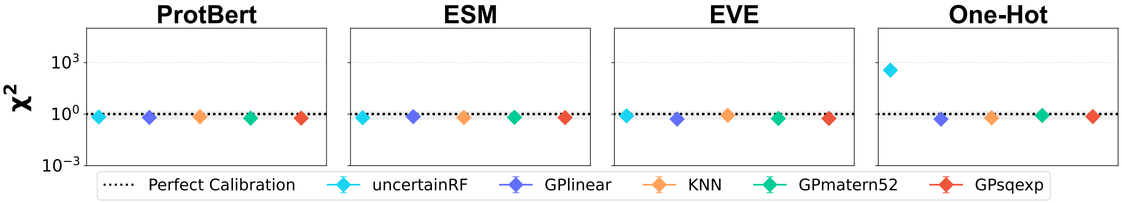


(c) $\chi^2$ calibration of regressors on TIM-Barrel when splitting by position.

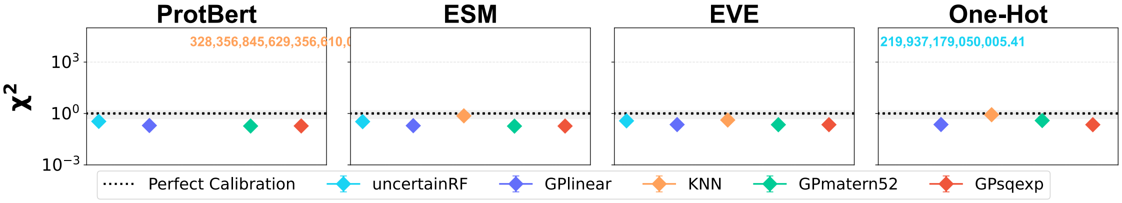


(d) $\chi^2$ calibration of regressors on T2-MTH when splitting by position.

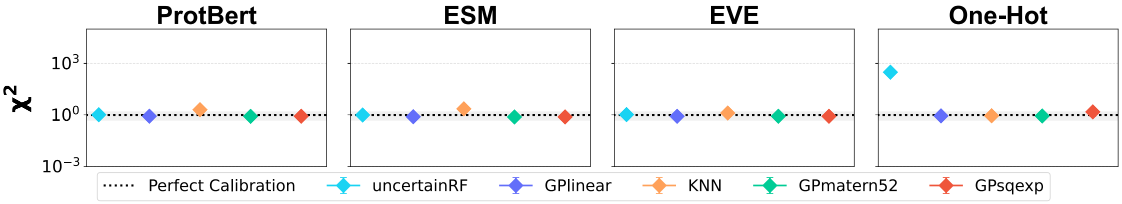


(e) $\chi^2$ calibration of regressors on BRCA1 when splitting by position.

Supplementary Figure 10: Calibration of regressors by (reduced) $\chi^2$ for splitting by position. In the case that the values are outside of the range (y-axis), the numerical value is displayed instead.

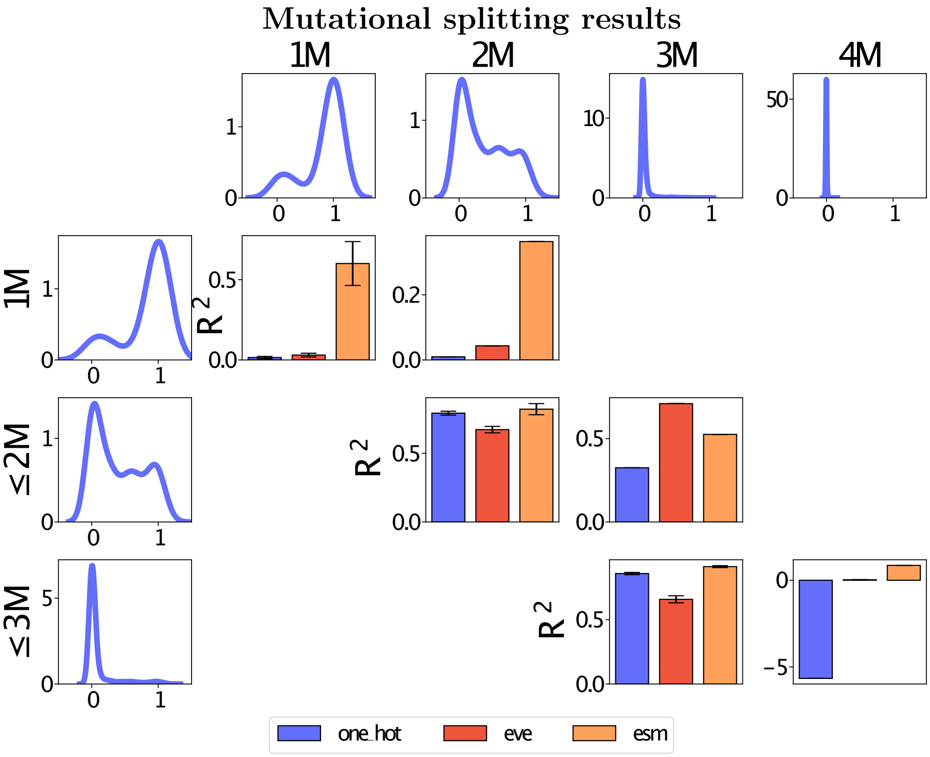


Supplementary Figure 11: $R^2$ metric on *mutation-degree* protocol on GP (Matern52) represented by ESM-1B. Functional observations of ParD-antitoxin are density curves in first row and column. We assess the adjusted $R^2$ score. The training domains are listed on rows and testing domains are columns, such that diagonals display in-domain performance, and off diagonal extrapolation performance. To assess in-domain performance a standard 5-fold CV protocol was used.

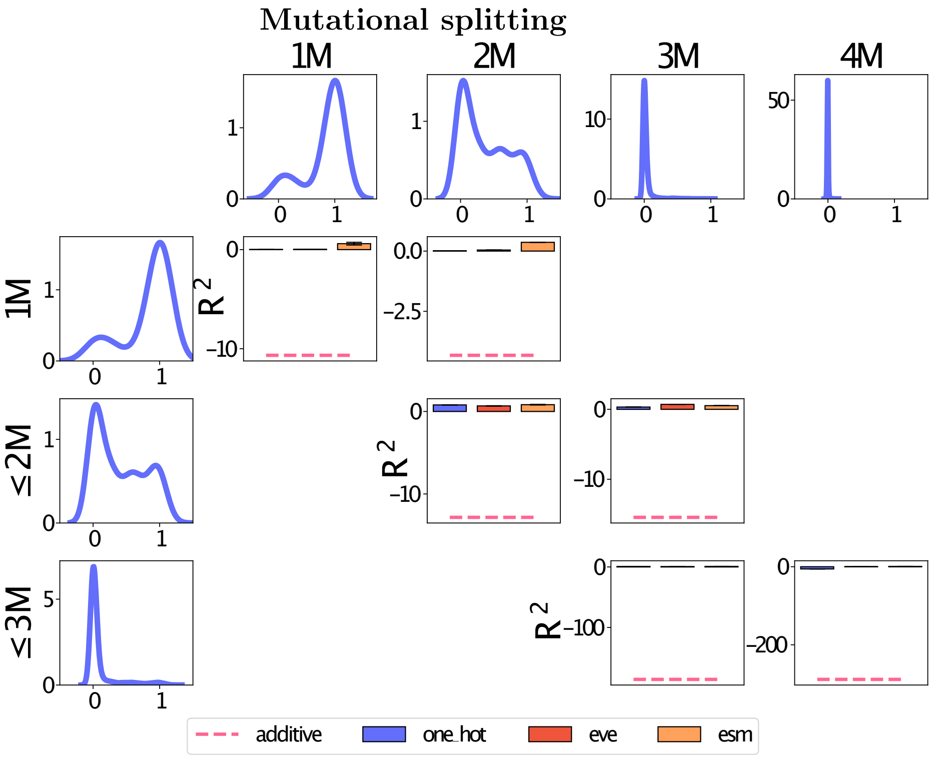


#### Results

Supplementary Figure 12: $R^2$ metric on *mutation-degree* protocol on GP (Matern52) represented by ESM-1B including *additive mutations* as reference baseline. Functional observations of PARD-ANTITOXIN are density curves in first row and column. We assess the adjusted $R^2$ score. The training domains are listed on rows and testing domains are columns, such that diagonals display in-domain performance, and off diagonal extrapolation performance. To assess in-domain performance a standard 5-fold CV protocol was used.

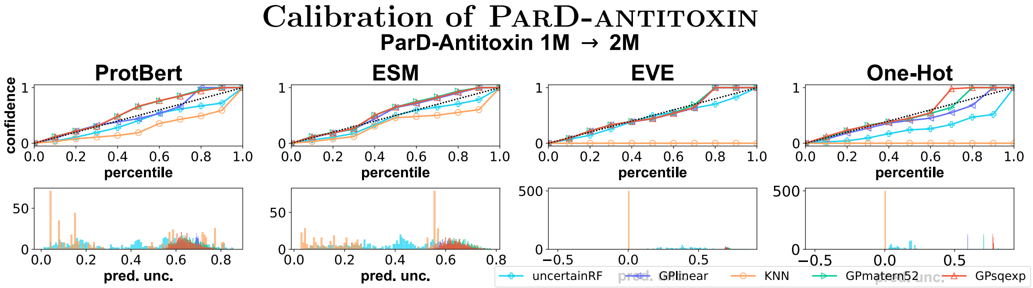


(a) Calibration of ParD-antitoxin (top) w.r.t. perfect calibration (dotted diagonal) and count of predictive variances (bottom) for the 1M to 2M extrapolation task.

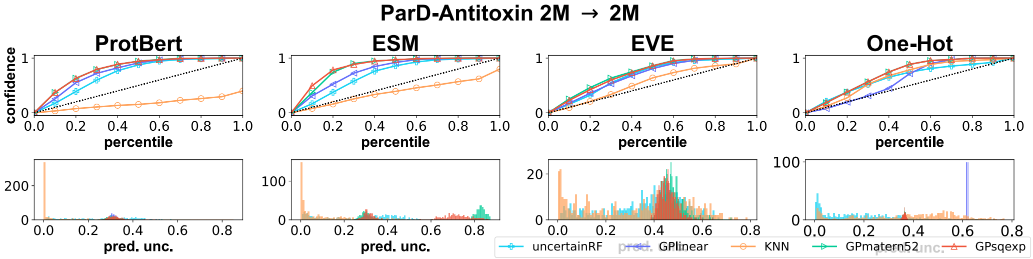


(b) Calibration of ParD-antitoxin (top) w.r.t. perfect calibration (dotted diagonal) and count of predictive variances (bottom) for the 2M to 2M intrapolation (5-fold CV) task.

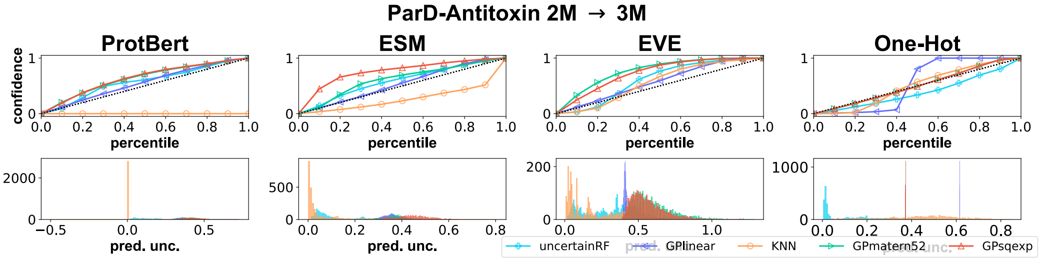


(c) Calibration of ParD-antitoxin (top) w.r.t. perfect calibration (dotted diagonal) and count of predictive variances (bottom) for the 2M to 3M extrapolation task.

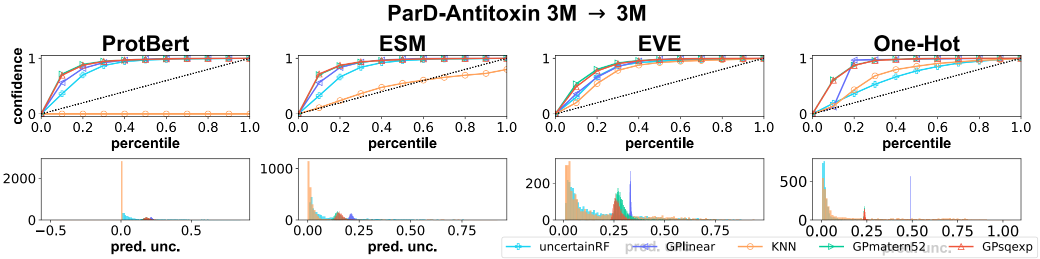


(d) Calibration of ParD-antitoxin (top) w.r.t. perfect calibration (dotted diagonal) and count of predictive variances (bottom) for the 3M to 3M intrapolation task.

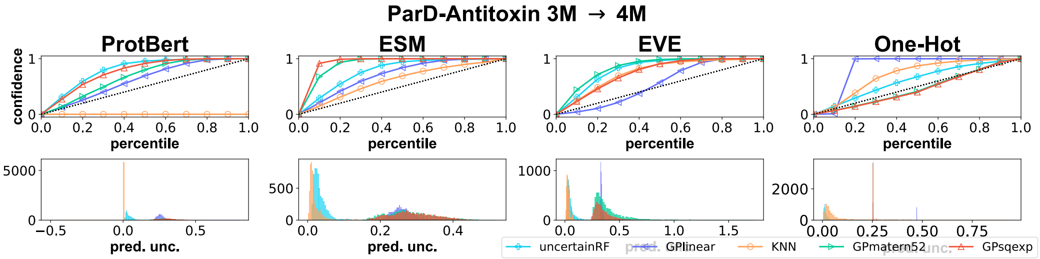


(e) Calibration of ParD-antitoxin (top) w.r.t. perfect calibration (dotted diagonal) and count of predictive variances (bottom) for the 3M to 4M extrapolation task.

Supplementary Figure 13: Calibration overview across extra- and intra-polation task assessments of ParD-antitoxin across representations (columns). Note the peaked predictive uncertainties (constant confidence curves) for KNN regressors with one neighbor predictions.

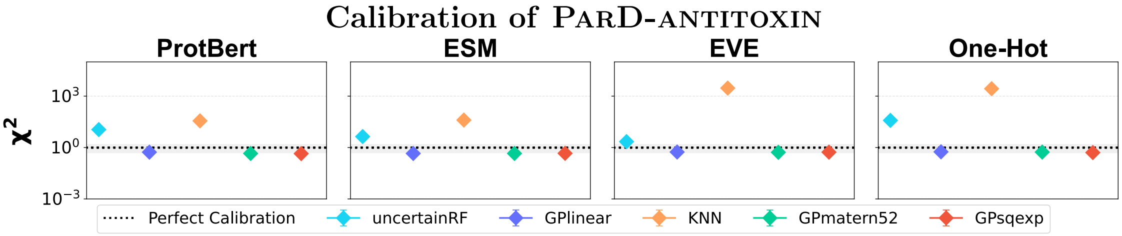


(a) Calibration by $\chi^2$ assessment of ParD-antitoxin (top) w.r.t. perfect calibration (dotted line and grey region) for the 1M to 2M extrapolation task.

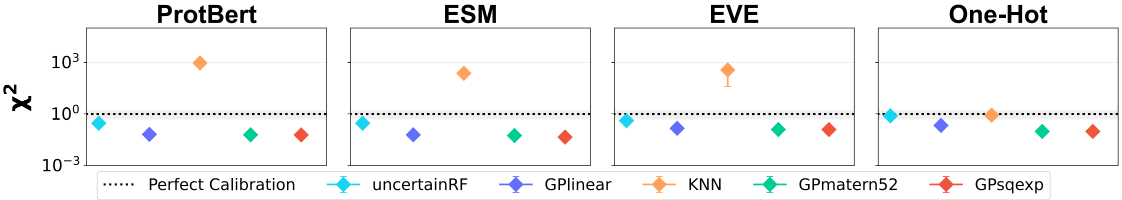


(b) Calibration by $\chi^2$ assessment of ParD-antitoxin (top) w.r.t. perfect calibration (dotted line and grey region) for the 2M to 2M intrapolation task.

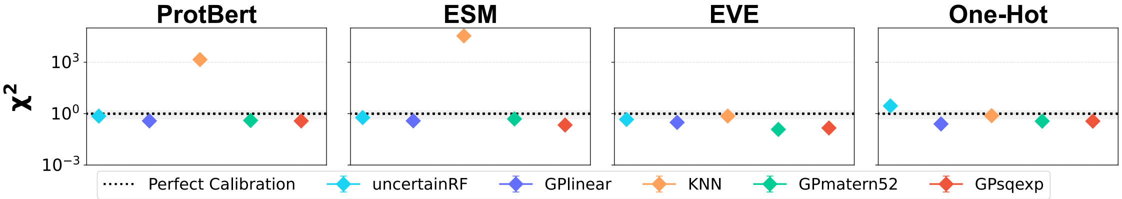


(c) Calibration by $\chi^2$ assessment of ParD-antitoxin (top) w.r.t. perfect calibration (dotted line and grey region) for the 2M to 3M extrapolation task.

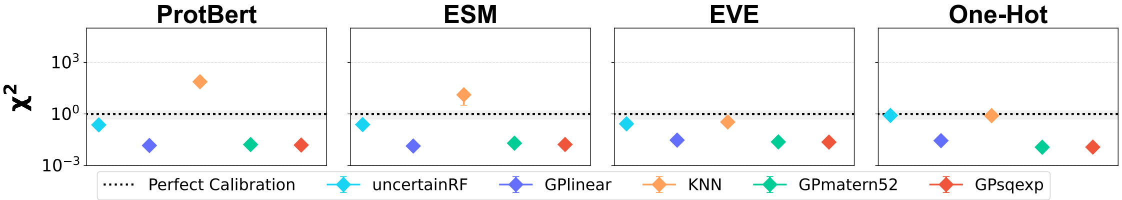


(d) Calibration by $\chi^2$ assessment of ParD-antitoxin (top) w.r.t. perfect calibration (dotted line and grey region) for the 3M to 3M intrapolation task.

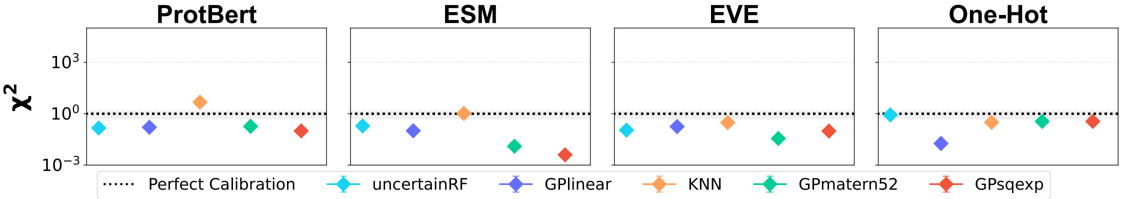


(e) Calibration by $\chi^2$ assessment of ParD-antitoxin (top) w.r.t. perfect calibration (dotted line and grey region) for the 3M to 4M extrapolation task.

Supplementary Figure 14: $\chi^2$ calibration overview across extra- and intrapolation task assessments of ParD-antitoxin across representations (columns).

### Optimization of $\beta$-Lactamase

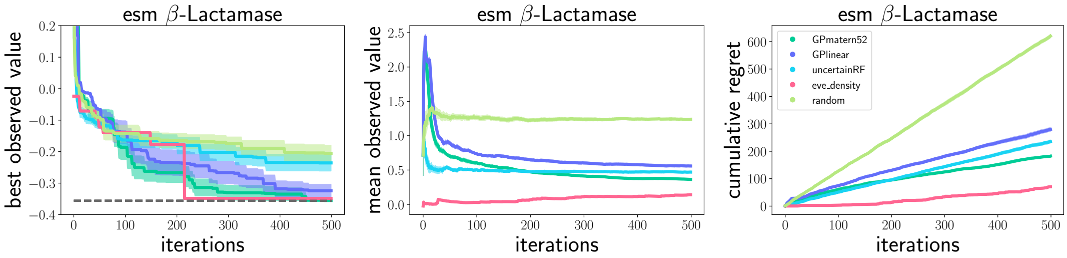


(a) Optimization experiment results of $\beta$-Lactamase on ESM-1B w.r.t. best observed value (left), mean value (center), and cumulative regret (right).

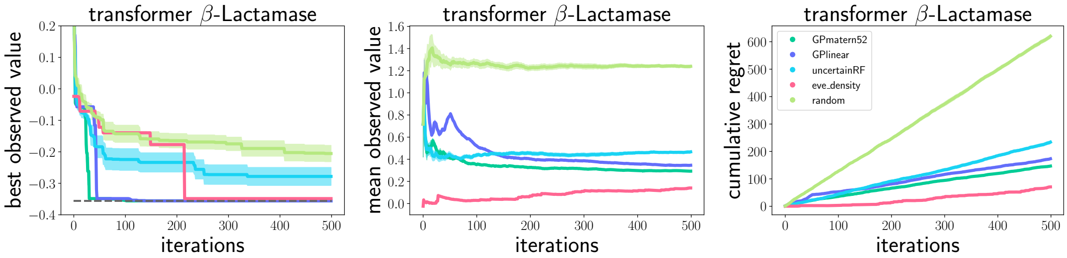


(b) Optimization experiment results of $\beta$-Lactamase on ProtBert w.r.t. best observed value (left), mean value (center), and cumulative regret (right).

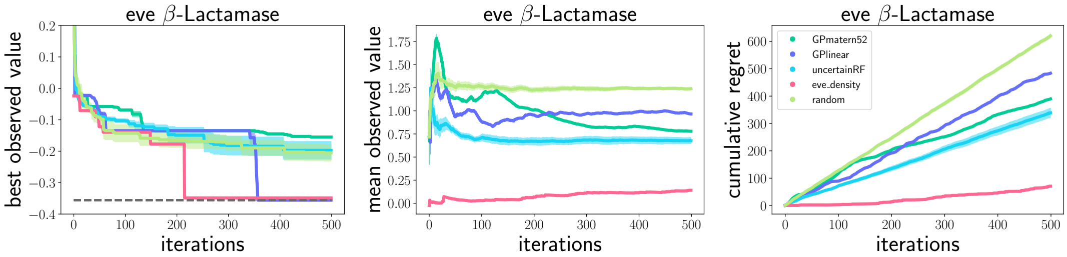


(c) Optimization experiment results of $\beta$-Lactamase on EVE w.r.t. best observed value (left), mean value (center), and cumulative regret (right).

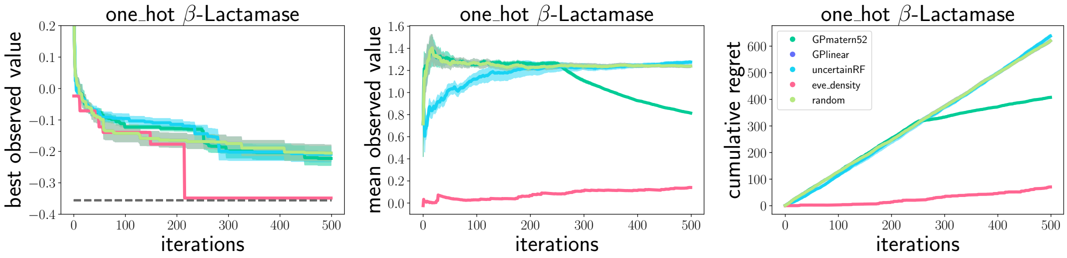


(d) Optimization experiment results of $\beta$-Lactamase on one-hot w.r.t. best observed value (left), mean value (center), and cumulative regret (right).

Supplementary Figure 15: Optimization as sequence selection with a budget of 500 steps. Across selected sequence candidates the best values (left), mean prediction (middle) and cumulative regret (right). For each method the mean prediction (bold line) from 10 different random seed runs, and standard error across (shaded regions). The reference baselines are randomly selecting observations (light green), and iterating over the ranked sequences by EVE scoring (pink). The best possible value in the setup is the dashed grey line (left), which is found by the GP models and scored ranking within the allotted budget.

#### Optimization of Ubiquitin

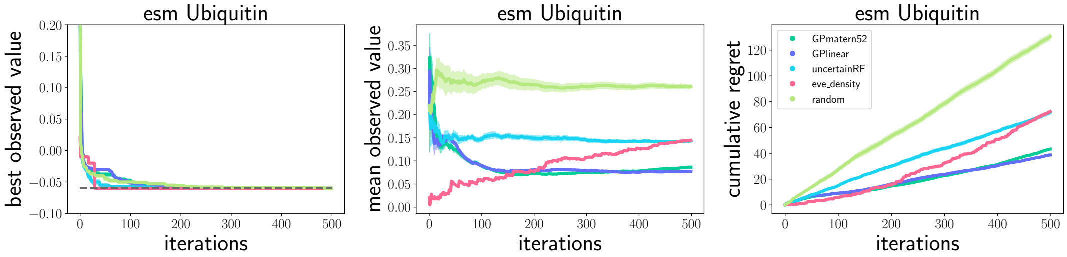


(a) Optimization experiment results of Ubiquitin on ESM-1b w.r.t. best observed value (left), mean value (center), and cumulative regret (right).

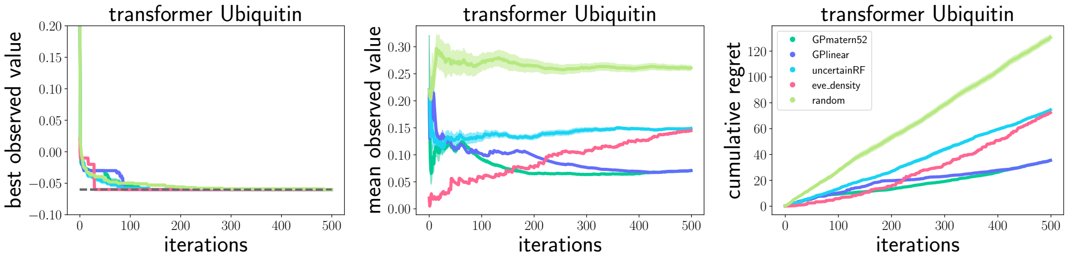


(b) Optimization experiment results of Ubiquitin on ProtBert w.r.t. best observed value (left), mean value (center), and cumulative regret (right).

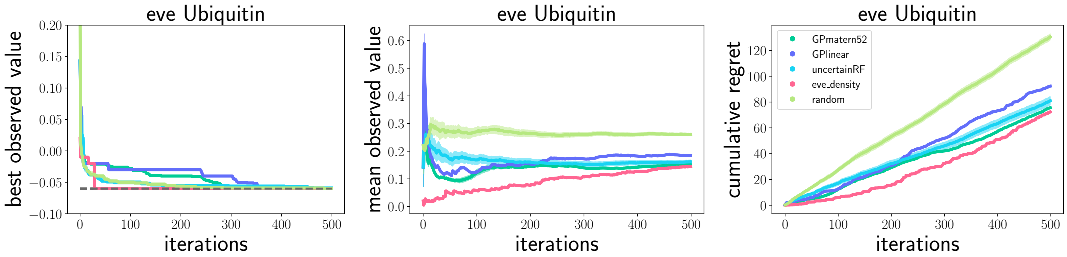


(c) Optimization experiment results of Ubiquitin on EVE w.r.t. best observed value (left), mean value (center), and cumulative regret (right).

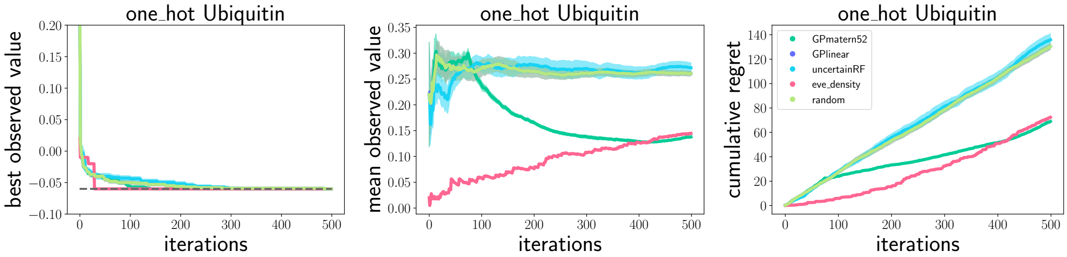


(d) Optimization experiment results of Ubiquitinon one-hot w.r.t. best observed value (left), mean value (center), and cumulative regret (right).

Supplementary Figure 16: Optimization as sequence selection with a budget of 500 steps. Across selected sequence candidates the best values (left), mean prediction (middle) and cumulative regret (right). For each method the mean prediction (bold line) from 10 different random seed runs, and standard error across (shaded regions). The reference baselines are randomly selecting observations (light green), and iterating over the ranked sequences by EVE scoring (pink). The best possible value in the setup is the dashed grey line (left), which is found by the GP models and scored ranking within the allotted budget.

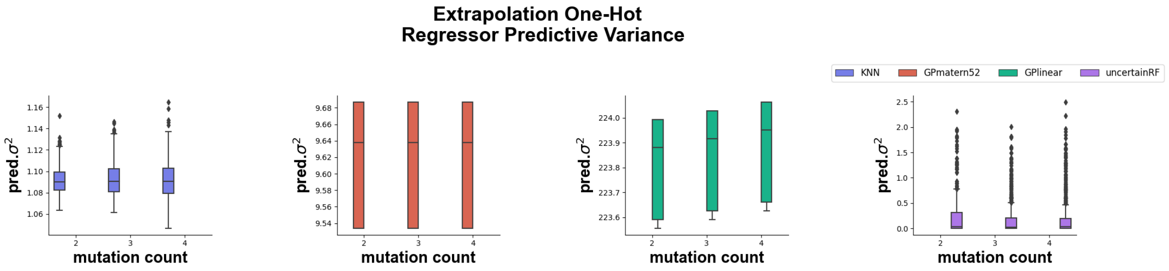


(a) Predictive variances of regressors, data represented via ONE-HOT

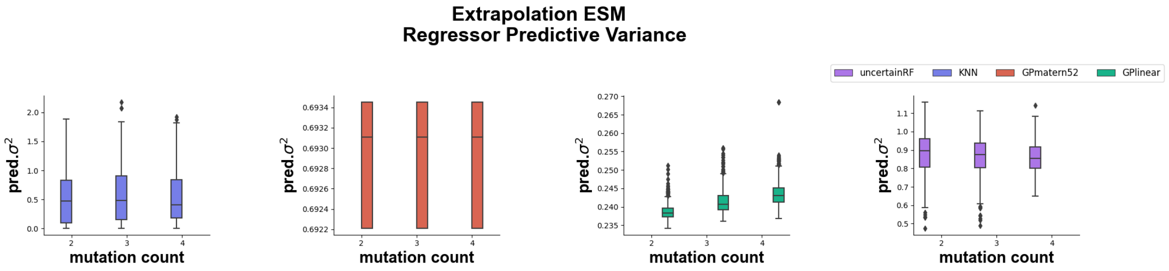


(b) Predictive variances of regressors, data represented via ESM-1B.

Supplementary Figure 17: Predictive variances of regressors tasked with extrapolation of randomly inserted mutations on the $\beta$-LACTAMASE dataset.

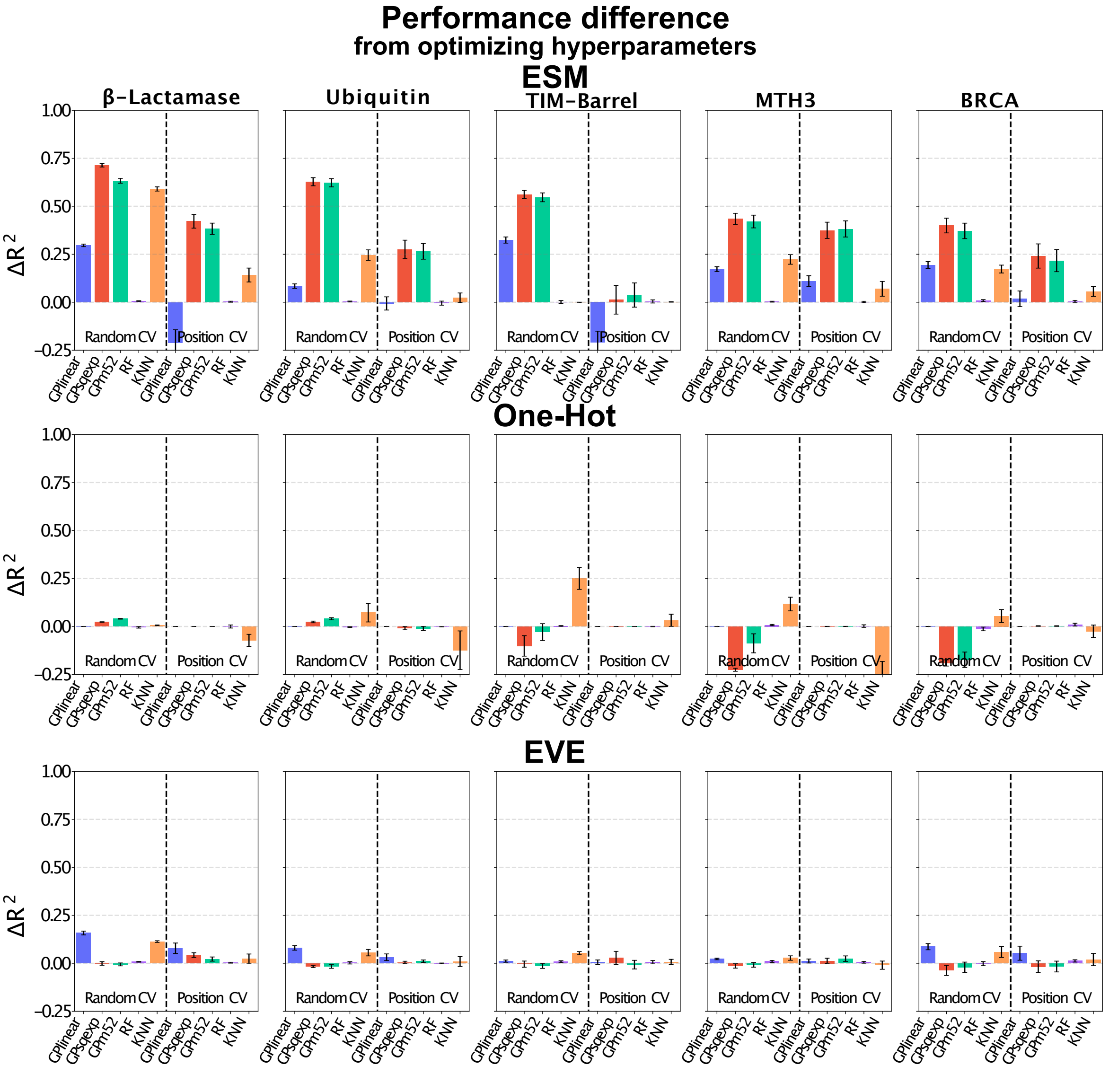


Supplementary Figure 18: Difference in performance ($R^2$) from parameter optimization for ESM-1B, ONE-HOT, EVEacross all regressors (x-axis).

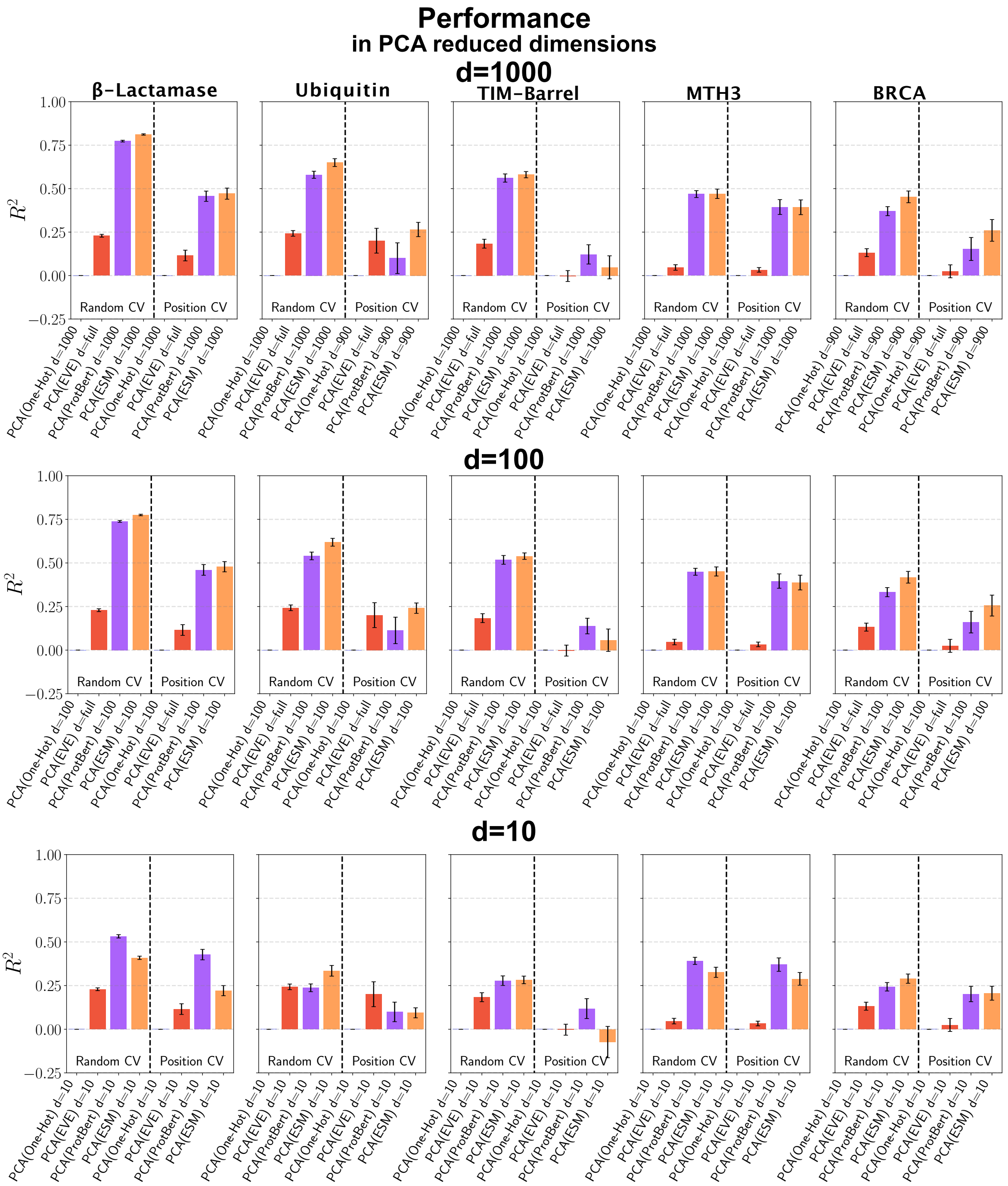


Supplementary Figure 19: Performance ($R^2$) of GPMatérn$\frac{5}{2}$ regressor on linearly dimensionality (PCA) reduced embeddings for random and positional splitting protocols. Note that for Ubiquitin not all $d = 1000$ representations were stable, hence the dimension has been reduced by an additional $\frac{1}{10}d$ so that $d = 900$, specifically in the positional splits. EVE dimensionality is less than 100, therefore d=10 is the only reduction - d=1000 and d=100 display the full EVE representation.

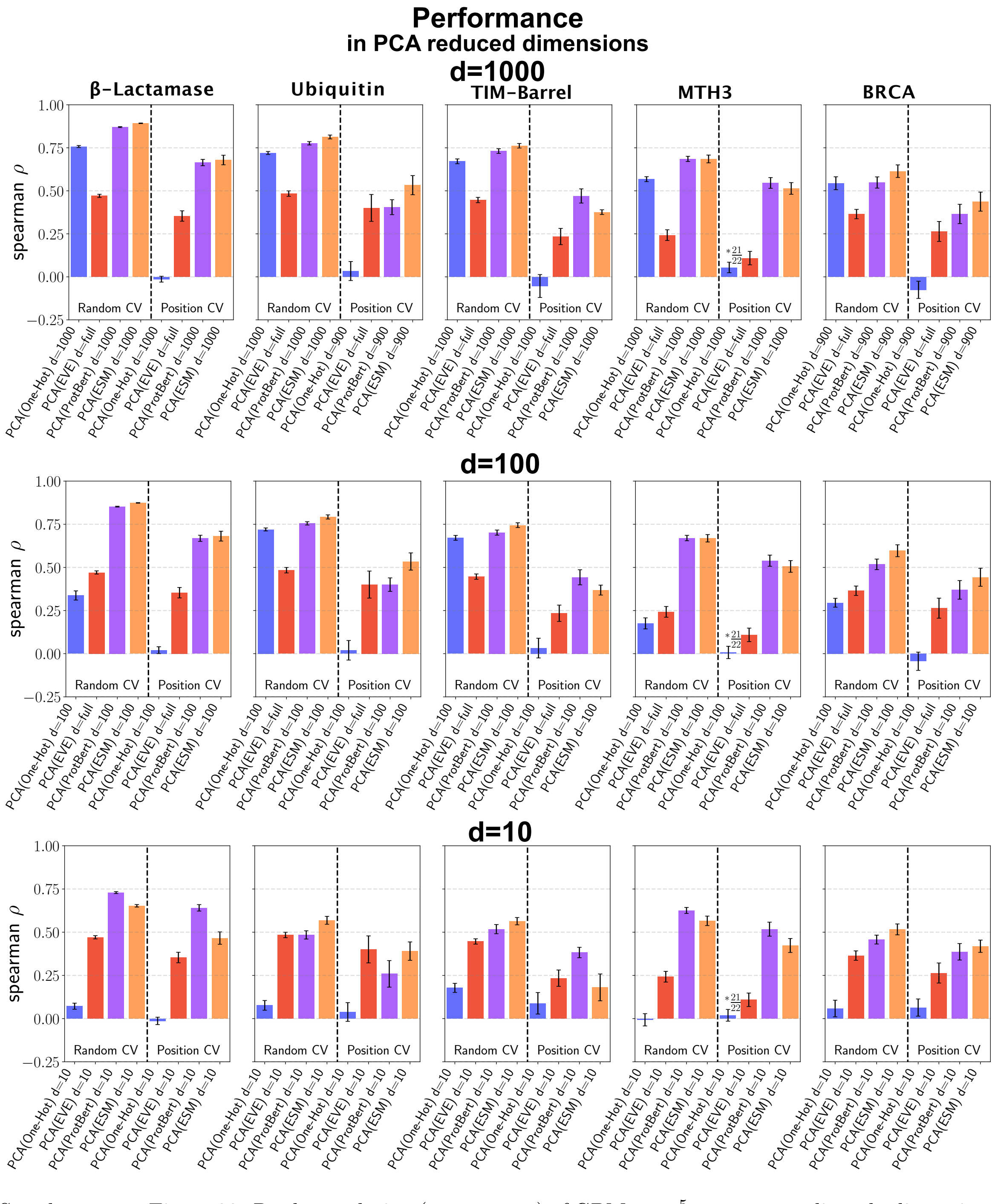


Supplementary Figure 20: Rank correlation (spearman $\rho$) of GPMatérn$\frac{5}{2}$ regressor on linearly dimensionality (PCA) reduced embeddings for random and positional splitting protocols. Note that for Ubiquitin not all $d = 1000$ representations were stable, hence the dimension has been reduced by an additional $\frac{1}{10}d$, s.t. $d = 900$, specifically in the positional splits. EVE dimensionality is less than 100, therefore d=10 is the only reduction - d=1000 and d=100 display the full EVE representation.
